# Role of combined cell membrane and wall mechanical properties regulated by polarity signals in cell budding

**DOI:** 10.1101/2020.04.30.071456

**Authors:** Kevin Tsai, Samuel Britton, Ali Nematbakhsh, Roya Zandi, Weitao Chen, Mark Alber

**Affiliations:** Department of Mathematics, University of California, Riverside, CA; Interdisciplinary Center for Quantitative Modeling in Biology, University of California, Riverside, CA; Department of Biochemistry and Molecular Biology, Penn State University, PA; Department of Physics, University of California, Riverside, CA

**Keywords:** yeast, budding, asymmetric cell growth, multi-scale, three-dimensional modeling, particle model, protein polarization, visco-elastic

## Abstract

The budding yeast, *Saccharomyces cerevisiae*, is a prime biological model to study mechanisms underlying asymmetric growth. Previous studies have shown that, prior to yeast bud emergence, polarization of a conserved small GTPase, Cdc42, must be established. Additionally, hydrolase changes the mechanical properties of the cell wall and plasma membrane with the periplasm between them (cell surface). However, how the surface mechanical properties in the emerging bud are different from the properties of the mother cell and their role in bud formation are not well understood. We hypothesize that the polarized chemical signal alters the local dimensionless ratio of stretching to bending stiffness of the cell surface of the emerging yeast bud. To test this hypothesis, a novel three-dimensional coarse-grained particle-based model has been developed which describes inhomogeneous mechanical properties of the cell surface. Model simulations suggest that regulation of the dimensionless ratio of stretching to bending stiffness of the cell surface is necessary to initiate bud formation. Furthermore, model simulations predict that bud shape depends strongly on the experimentally observed molecular distribution of the polarized signaling molecule Cdc42, while the neck shape of the emerging bud is strongly impacted by the properties of the chitin and septin ring. This 3D model of asymmetric cell growth can also be used for studying viral budding and other vegetative reproduction processes performed via budding.

## 1 Introduction

In this paper, a model is introduced for studying asymmetric cell growth, a prominent reproductive process utilized in many organisms to generate cell diversity during development. It is also important in determining cell fate when, for example, stem cells divide for the purpose of proliferation or differentiation. The budding yeast *Saccharomyces cerevisiae* is a fungus that can reproduce via asymmetric growth and serves as a classic model to study the principles underlying this fundamental process [1–8]. Reproduction in yeast, i.e. budding, is a delicate process governed by a combination of dynamically changing biochemical signaling networks, turgor pressure, transport of subcellular organelles, and regulation of the mechanical properties of the yeast cell surface, consisting of cell wall, cell membrane and periplasm between them.

Structurally, the yeast cell wall is a dynamic network primarily composed of polysaccharides. The cell wall network is composed of 1,3- β -glucan, 1,6- β -glucan, and a relatively small amount of chitin proteins. Linkage between the 1,3- β -glucan, 1,6- β -glucan, and chitin proteins are established to maintain the structural integrity of the cell wall [10]. Beneath the cell wall structure lies the cell membrane consisting of lipids and membrane-bound proteins similar to the membrane in animals. While the cell wall mechanically supports the cell integrity in response to forces from the environment and maintains cell shape, the cell membrane acts as the barrier to the free diffusion in the cytosol, provides binding sites for molecular signaling pathways involved in the biosynthesis of cellular components, and relays the environmental conditions to the cell interior via signaling transduction pathways to regulate the osmotic balance [11].

Yeast budding starts with a protrusion in the cell surface and results in cell division to form a daughter cell separated from the mother (**Figure 1A**). A single bud is generated in one cell cycle. Notice that no nucleus is formed yet in the bud at the early stage (**Figure 1B**) [12]. The location of the protrusion site, or the bud site, is determined by asymmetric distribution of Cdc42 and growth-associated proteins established before cell shape change, which is followed by a polarization of structural components including actin cables, septin, and myosin [13,14]. These polarization events play an important role in budding. It has been shown experimentally that multiple concurrent protrusions during the budding process occur when the Cdc42 signaling pathway is impaired [15].

**Figure 1.**
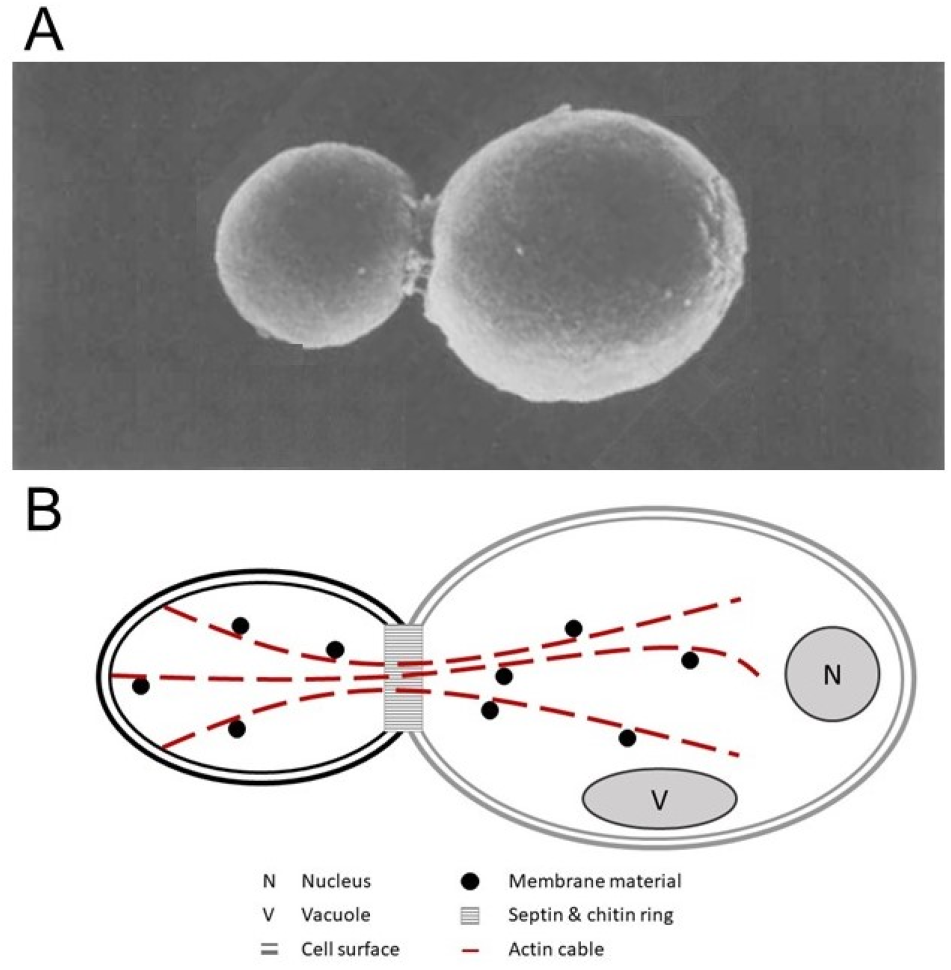
Experimental image (A) and representative diagram (B) of the yeast mother cell (right) and the developing bud (left) separated by the chitin and septin ring. (B) Cell wall (outer boundary) and membrane (inner boundary) are represented by two curves. Internal components of the mother cell include nucleus and vacuole. Actin cables (dashed red lines) polarize at the bud site and recruit new cell membrane/wall materials (black points). (Image A is reproduced with permission from Hanschke et al. [9]).

Shortly before the protrusion occurs, septins and chitins within the cell membrane and cell wall are assembled to form ring-like structures [16,17] (**Figure 1**). It has been shown that the septin ring and the chitin ring, located in the cell membrane and cell wall, respectively, have similar functionality in controlling the size of the budding neck via different mechanisms, and the synthases responsible for the assembly of these two rings are related [18]. Presence of the rings is essential. They colocalize and, along with the linkage between the chitin ring and 1,3- β -glucan, limit the expansion along the neck during budding. Moreover, these two rings, especially the septin ring, act as a diffusion barrier impacting bud morphogenesis [6,19]. Meanwhile, actin cables polarize to direct the transport of secretory vesicles as well as new cell membrane and cell wall materials from the cytosol to the budding site. Several studies have suggested that mutants which have improper formation of the chitin and septin rings or polarized actin cables give rise to wide budding necks, which can be detrimental to the survival of the cell [20–22].

During the early stages of the budding process, the mother cell exhibits marginal change in size and the turgor pressure remains sufficiently constant in the rage of 0.1 – 1.0 MPa [23–25]. A recent study also suggests that during the entire reproduction cycle, the turgor pressure remains at approximately 0.21 MPa [26]. While the turgor pressure acts as the driving force to generate a bulge on the cell surface, Cdc42 polarizes and regulates cell surface mechanical properties to maintain cell integrity. This is accomplished by degrading the chemical bonds between polymers within the 1,3- β -glucan network in the cell wall and recruiting new materials to the cell wall and cell membrane [14]. However, it is challenging to experimentally measure mechanical properties of the cell surface in actively growing cells. Recently, the elasticity of the cell wall was measured during the yeast budding process and it was found that stiffness of the bud was slightly higher than that of the mother cell, although the obtained value may depend on the timing of measurements and the highly curved surface [27]. This result is different from an earlier observation in which the cell wall at the budding site becomes less rigid prior to bud emergence [14,28]. It, therefore, remains unclear whether the change in mechanical properties of the yeast cell surface is necessary for the bud emergence. Moreover, it is not known how the mechanical properties of the cell surface are regulated to form a bud with appropriate shape.

Multiple computational models have been developed to propose and test different mechanisms underlying the budding process such as the clathrin-mediated endocytosis [29,30] and the wrapping of nanoparticles [31–33]. In Gompper et al. [34–36], a tether-and-bead model was developed to study cell surfaces with fixed sizes and fluctuating topologies, where the cell surface was discretized by triangulated mesh and a probabilistic re-meshing algorithm was introduced to represent cell growth. This model has been extended into a particle-based framework to study the effect of molecular turnover and the material exchange between the cell membrane and cytosol on the cell shape by incorporating a mesh-refinement algorithm [37] to facilitate the stability of the model in capturing local cell surface deformation.

On the other hand, several computational models have been developed to study morphogenesis based on description of the entire inhomogeneous cell wall. For example, in the model for mating yeast, cell wall was described as an inhomogeneous viscous fluid shell and coupled with the cell wall integrity signaling pathway, which governs the wall synthesis and controls its stiffness, to study the coordination of mechanical feedback in cell wall expansion and assembly in mating yeast [38,39]. Similar approach has been applied to study the tip growth of the pollen tube in plants [38,39]. In a paper [27,40], the interplay between the turgor pressure and the elasto-plasticity of the cell wall during yeast budding was investigated by using a single-cell growth model (SCGM). Mother cell and bud are represented as two separate spheres with identical wall elasticity but different levels of plasticity. Growth is described by the dynamics of cell radius, which is impacted by different levels of wall plasticity and fluctuating turgor pressure. Simulation results suggest that the bud must be significantly more exposed to plastic expansion compared to the mother cell in order for proper bud formation to take place.

In this paper, a novel 3D coarse-grained particle-based model is described and used to examine the impact of changing mechanical properties of combined cell membrane and cell wall (called cell surface hereafter) on the bud formation in yeast cells. Specifically, model simulations show how local cell surface growth and deformation in an early bud formation, controlled by experimentally observed polarized distribution of Cdc42, impacts the global deformation of the cell surface. Model parameters were calibrated to resemble the Young’s modulus of the cell wall measured in experiments [41]. The model assumes that the turgor pressure remains constant throughout the early stages of the budding process which we focus on in this paper. Additionally, the model assumes ratios of stretching to bending modulus of the bud and mother cell to be different from each other, which is based on the experimentally observed presence of wall-degrading enzymes at the bud site. (Description of the terminology used in the paper is provided in the **SI Table 4**).

Model simulation results indicate that increased dimensionless stretching to bending stiffness ratio, Föppl-von-Kármán number, within the bud region at the early stage is essential for the bud emergence and maintenance of the spherical shape. Chitin and septin rings were shown to impact the neck shape without changing the bud sphericity, as well as reducing the high Föppl-von-Kármán number required for bud emergence. By varying the distribution of the polarized mechanical regulator, it was demonstrated that reduced polarized signal distribution leads to asymmetric bud formation. Moreover, by assuming that the mechanical properties at the bud site can recover in time to the same level as those of the mother cell, it was shown that buds can acquire a symmetric shape similar to those observed in experiments. The new model can be extended to study the impact of dynamical changes of molecular distributions in yeast budding, as well as viral budding and other vegetative reproduction processes performed via budding.

## 2 Methods

### 2.1 General model description

Yeast mother cells can be either egg-shaped, elliptical, or spherical [42]. While the size of an yeast cell varies depending on the cell types, for simplicity we assume that mother cell initially is a sphere with radius of 2.0 μ*m* [43]. The sphere, representing the cell wall and membrane, is discretized into a triangulated mesh. The triangulated surface is a simplified representation of the elastic network of the yeast cell surface. The nodes are connected by linear springs in each triangle to capture the in-plane elasticity, whereas the bending springs are applied to triangles sharing a common edge to model the out-of-plane elasticity (**Figure 2**) [44–46]. This mesh discretization allows calibration of the model elasticity using experimental data by probing the elastic response under stress (Section 2.5). We also assume that the bud site has been predetermined and Cdc42 becomes polarized to regulate the elasticity at the bud site.

**Figure 2.**
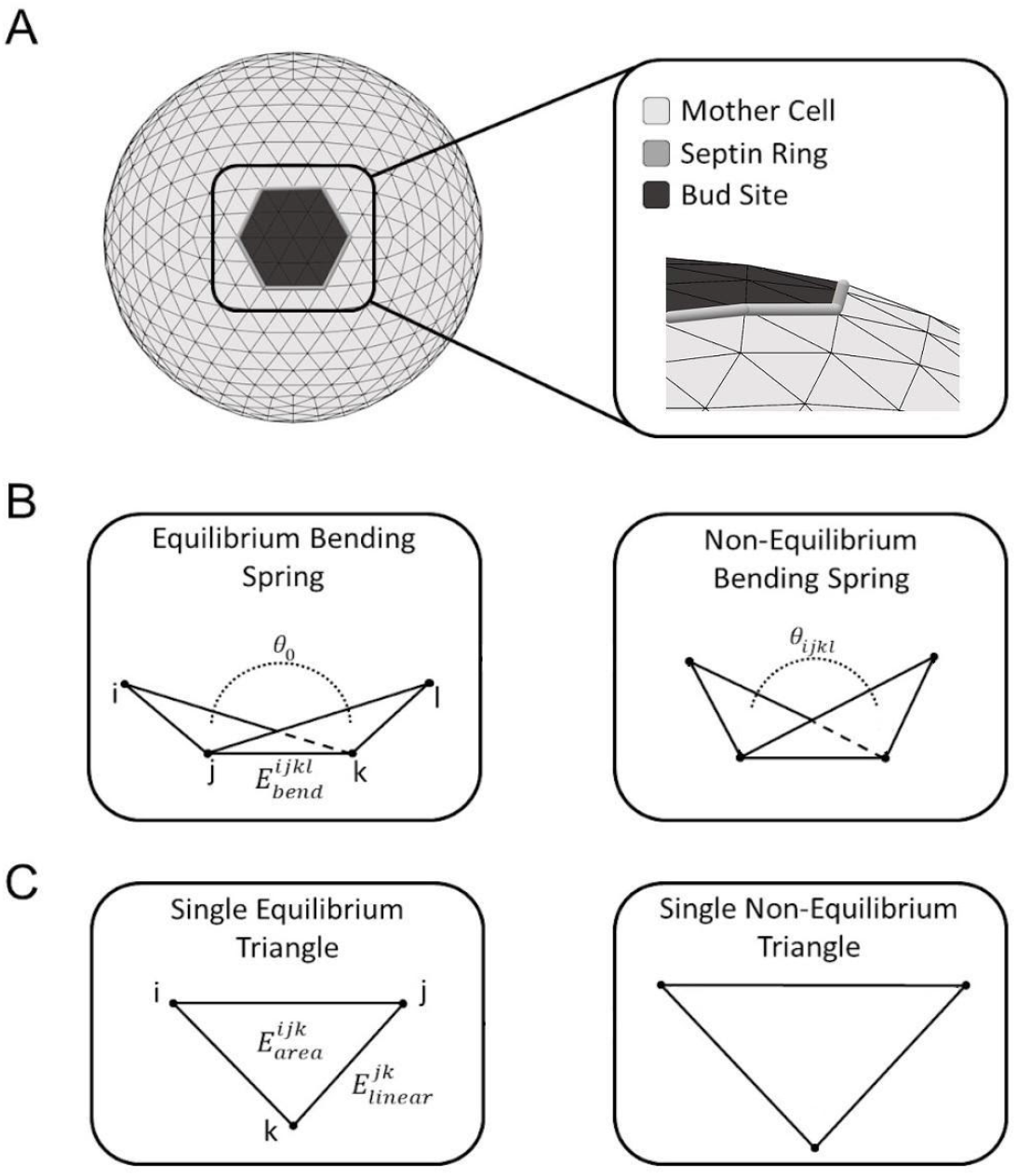
Schematic diagram of components comprising the *in silico* yeast cell model. (A) Initial simulated mother cell representation including predetermined bud region (dark grey), combined chitin and septin rings (grey tubes), and mother cell surface (light grey). (B-C) Individual model spring elements are shown at equilibrium (left) and non-equilibrium (right).

The bud site is enclosed by the chitin and septin ring which is approximately 1 μ*m* in diameter [44]. The size of the ring remains unchanged throughout the budding process. While the mechanical role of the ring has yet to be confirmed experimentally, the lack of the change of the ring and bud neck size, indicates a constraining effect. In the model, the bud neck is represented by using a set of linear springs with a stiffness an order of magnitude higher than the one used to model the surface of the mother resulting in a rigid ring type behaviour. Furthermore, the ring acts as a demarcation separating bud site and mother cell (**Figure 2**) [44,47]. Distinct mechanical properties are assigned to the bud surface under the influence of wall degrading enzymes and to the mother cell surface. Previously developed re-meshing techniques [34,35,37,48,49] are utilized in the model to capture the structural response to osmotic pressure [50,51].

#### Computational implementation

The iFEM [52], a MATLAB software package, was used to generate the initial mesh configuration. The density of the triangulated mesh can be changed according to the requirement of the resolution of the modeling system. In all simulations included in this paper without specification, the number of triangles used for the mother cell at the beginning of the simulation is 1280, and out of them 24 triangles belong to the budding region.

### 2.2 Equations of motion

Motion of each node *i* from the model representation of the cell surface is described by the following equation:

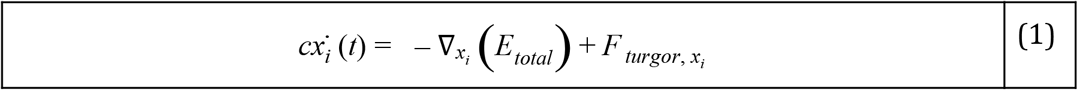

where *c* is the friction coefficient depicting the viscosity of the cell surface, represents the gradient with respect to the *i^th^* node position, *E_total_* represents the total potential energy used to model the mechanical properties of the surface of the cell and *F_turgor_* is the force derived from the force-stress relationship originating from the constant turgor pressure acting on the cell surface. (More details are provided in Sections 2.3-2.4.)

We define the total potential energies used to model the cell surface: *E_total_ = E_linear_ + E_area_ + E_bend_ + E_volex_*. Here *E_linear_* is the linear potential between connected nodes and *E_volex_* is the Morse potential describing interactions between nodes that are not connected by an edge, modeling the self-avoiding property preventing cell-cell intersection. *E_area_* and *E_bend_* represent area and bending potentials yielding forces that control the area expansion resistance of each individual triangular element and level of bending between triangular elements. Due to the micron scale of the yeast cell and fluid environment required for yeast reproduction, the surrounding microenvironment acts as an overdamping media. We therefore assume that the cell in the model is in the overdamped regime where the inertia force is negligible [37,53]. Between two successive re-meshing steps, we simulate *N_relaxation_* = 200 iterative steps. *N_relaxation_* is chosen such that the relative change in total energy of the system converges to a value below a threshold of 0.01.

Notice that each mechanical potential comprising *E_total_* is chosen to represent cell surface elasticity, shape maintenance, and surface incompressibility, as described in Section 2.3. While the choice of the components of energy potential function is not unique, we believe that our results is inline with other form of energy potential function which have previously been used to describe the cell surface [37,54–56]. Potentials in the model were calibrated via simulated cell stretching tests, and adjusted to match the experimental atomic force microscopy (AFM) data, as described in Section 2.5.

### 2.3 Interaction potentials

We assume linear stiffness of the cell surface under low stress and use the internodal potential for nodes connected via an edge in the following form:

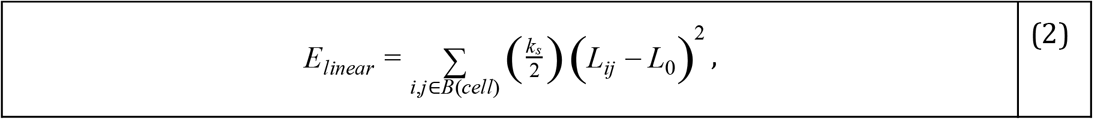

where *k_s_* is the linear spring coefficient, *L_ij_* is the length of the spring connecting node *i* and node *j*, and *L*_0_ is the equilibrium length of the bond. The sum is taken over all edges of the mesh, denoted by *B*(*cell*).

Following previous work [37,54], the local area expansion resistance of each individual triangular facet is represented using a harmonic potential:

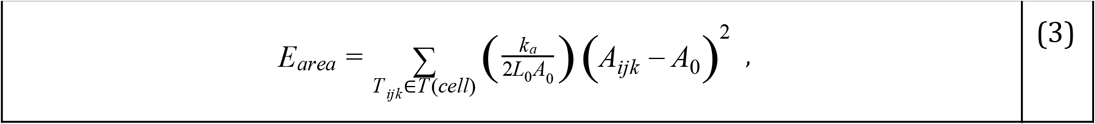

where *k_a_* is the area expansion resistance coefficient, *A_ijk_* is the current area of the triangle *T_ijk_, A*_0_ is the equilibrium triangle area, and *L*_0_ represents the equilibrium edge length in each triangular element. The sum is taken over all triangle elements *T*(*cell*).

To maintain the spherical shape, we adopt the approach that utilizes the angle-bending potentials between neighboring triangles that share a common edge to enforce cell curvature. In particular, we apply the method proposed in [57]. The explicit relationship between the equilibrium angle and the radius of curvature is 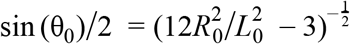 where θ_0_ is the equilibrium dihedral angle between unit normal vectors of edge-sharing triangles, *R*_0_ is the radius of curvature, and *L*_0_ is the equilibrium edge length of the triangle. Hence, the bending behavior is determined by the cosine bending potential based on the unit normal vectors of edge-sharing triangles:

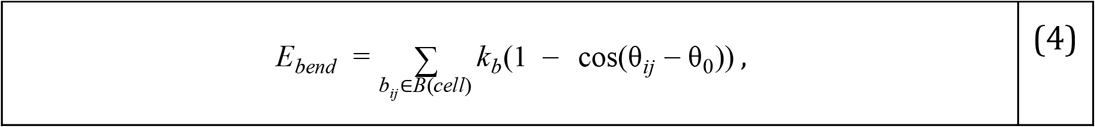

where *k_b_* is the bending coefficient, θ_*ij*_ is the current dihedral angle between the unit normal vectors of two triangles sharing edge *b_ij_*, and θ_0_ is the equilibrium dihedral angle. The sum is taken over all edges over the surface *B*(*cell*).

Volume exclusion constraints, *E_volex_*, are introduced to incorporate the self-avoiding property of different domains of the cell surface avoiding each other. Several models, whether with or without defining an absolute minimum distance between cell surface nodes, have employed compression-resistance potential [37,54]. The self-avoiding property is also necessary to maintain the numerical stability and cell surface topology to avoid concavity. We apply the standard Morse potential to enforce the self-avoiding property for non-connected nodes:

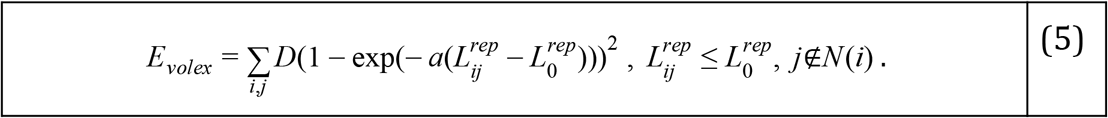

Here *D* represents the well depth of the Morse potential with width *a*. 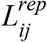 represents the distance between any two non-connecting nodes, 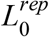 is the optimal distance between any two non-connecting nodes, and *N*(*i*) is the collection of nodes connected to a given node *i*. Parameter values and the calibration are described in Section 2.5.

### 2.4 Modeling cell growth

During the budding process, the yeast cell undergoes a local deformation with a narrow neck formation and cell surface material insertion to the bud site. To capture both geometric change and the expansion of the surface, we incorporate a re-meshing technique into the model.

To alter the edge-connectivity in the current mesh, which is to obtain a new geometry favoring bud formation, we employ the Monte Carlo edge re-connectivity algorithm. The Monte Carlo simulation for edge re-connectivity (also termed bond-flip) is a well-established approach and has been shown to effectively capture the topological change which is essential in surface deformation and protein folding [35,48,49]. The edge re-connectivity is determined probabilistically via energy-based comparison between the pre- and post-flip of edges in the existing mesh.

In order to describe an increase of the area of the bud, we define a quantity called strain associated with the area expansion

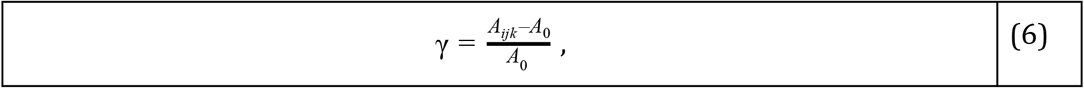

where *A*_0_ is the equilibrium area and *A_ijk_* is the current area of the triangle. During the simulation, if the value of γ exceeds a critical value, 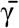, then new triangles are introduced into the system following the approach from [37]. Physically, this corresponds to the instance when new material insertion to the cell surface from the bulk is more energetically favorable. Biologically, it represents the response of a cell to excessive mechanical stress. Specifically, if the relative change in average area of two adjacent triangles, *T*_1_ and *T*_2_, is larger than 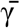, a new node, *m*, is introduced at the center of the shared edge (**Figure 3**). *T*_1_ and *T*_2_ are subsequently divided using the newly placed node, thereby creating four new triangles, *Q*_1_,…, *Q*_4_.

**Figure 3.**
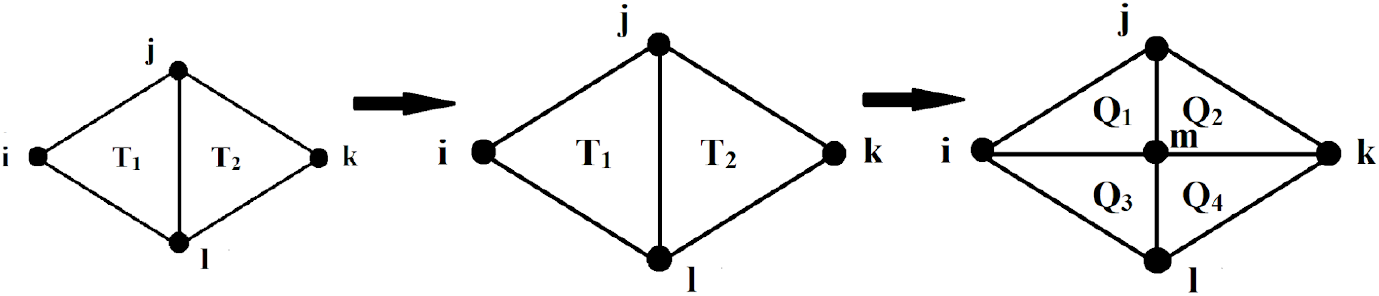
The growth (expansion) algorithm. Initial pair of triangles with a common edge (left) expands under stress (middle) resulting in a triangulation (right) after addition of a new node and new edges. This algorithm is used if the average area of *T*_1_ and *T*_2_ exceeds the critical value of γ.

We utilize this growth algorithm after *N* edge re-connectivity attempts described at the beginning of this section. Unless otherwise specified, *N* = 100. We check this growth condition over every pair of triangles sharing a common edge inside the budding region, and call this sweeping process as one growth step.

We assume that the turgor pressure is constant at the initiation of the budding stage we model, and the surface deformation at every step is small. For each triangle in the model, the force due to the turgor pressure acting on the node *i* is as follows,

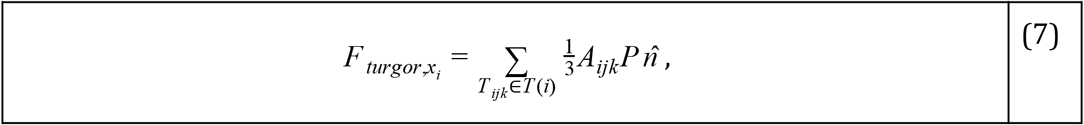

where *A_ijk_* is the current area of the triangle *T_ijk_*, *P* is the constant turgor pressure and 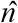 is the outward unit normal vector of the triangle. The sum is taken over triangles containing node *i*, denoted by *T*(*i*), in order to obtain the consistent force due to the turgor pressure applied to the overall cell surface [26].

### 2.5 Model calibration

In experiments, whole cell compression [58] and probing via atomic force microscopy (AFM) [41] are common approaches to measure the mechanical properties of a single cell including the stretching modulus and bending modulus on the cell surface.

While both methods have been applied to identify the elasticity of the yeast cell wall, the reported values differ from each other by up to two orders of magnitude. The model described in this paper was calibrated by using the measurement of the cell surface elasticity obtained by AFM in [41]. In this experiment, a nanoindentation on the cell wall was created and the modulus of elasticity was identified as 1.62 ± 0.22 MPa for a yeast cell prior to bud emergence.

During calibration, linear stiffness, bending stiffness and area expansion resistance coefficient are calculated after applying forces of the same magnitude in the opposite directions to a single cell having initially a spherical shape (**Figure 4A**). Re-meshing is not allowed in this simulation by assuming that the cell is not actively growing at this step of the algorithm. To reduce the sample size required in exploring the parameter space, we apply Latin Hypercube Sampling to generate a sufficient amount of samples distributed over the wide range [59,60]. Model simulations with the parameter set *k_s_* = 2.0, *k_a_* = 2.0, *k_b_* = 0.5 obtained as a result of calibration, produced an elastic response which results in the correct material behavior of the cell surface, plotted in a dashed line, falling within the experimental data shown as solid lines in **Figure 4B**. We use this parameter set as the wild type condition for the mother cell in the following sections unless specified otherwise. Parameter values used in the model simulations, including the equilibrium edge length and dihedral angle, are provided in the supplemental information (**Table S1**).

**Figure 4.**
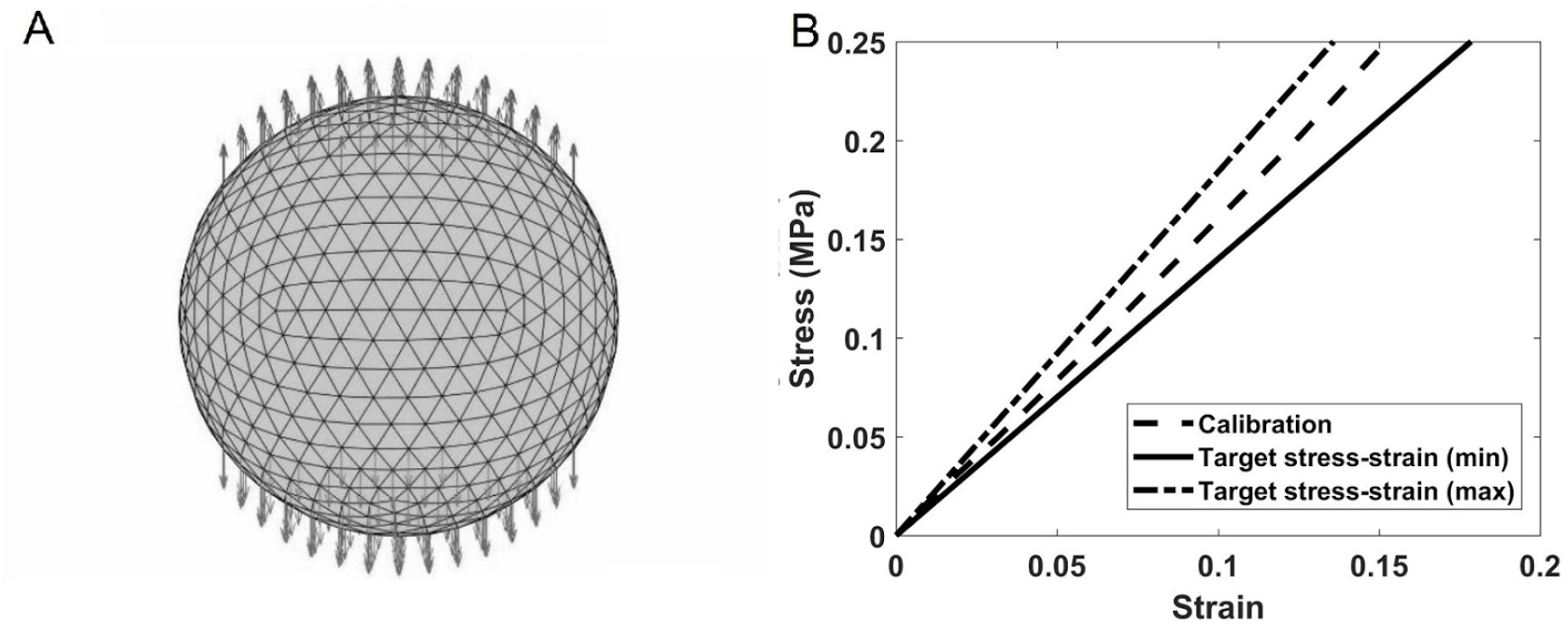
Calibration of the mechanical model for a single yeast cell without budding. (A) Forces of opposite directions are applied to detect the elasticity properties of the cell based on a chosen parameter set. (B) The stress-strain ratio (dashed line) using parameters *k_s_* = 2.0, *k_a_* = 2.0 and *k_b_* = 0.5.

## 3 Results

Prior to budding, Cdc42, small molecule GTPase, forms a cluster at a predetermined cortical site and orients actin cables toward the cluster. These cables then direct delivery of more Cdc42 as well as new cell membrane and cell wall materials using secretory vesicles to the cluster. This establishes a positive feedback loop to help establish the Cdc42 polarization and regulates cell surface mechanics at the bud site to prepare for budding [14,61,62].

The 3D computational model described in the previous section is used to determine whether changes in the cell surface elasticity at the bud site is required for bud formation and how these changes impact bud shape. In particular, we tested time-independent, spatial-dependent, and time-dependent changes in the mechanical properties, based on the spatiotemporal distribution of Cdc42 observed at different stages in experiments.

To model the impact of Cdc42 on the bud site enclosed by chitin and septin rings, each coefficient *k*_*_ defined in Eq. 2-4, is multiplied by a weight constant α_*_ varied between 0.0 and 1.0 (**Figure 2**). α_*s*_, α_*b*_, and α_*a*_, denote the weights for the stretching coefficient, bending coefficient, and the area expansion resistance coefficient, respectively.

In each simulation, the maximum number of iterative steps is *N* = 1.6×10^7^ with a total number of 800 growth steps. Since the focus of the paper is on the early stage of the budding process, a simulation is terminated when all growth steps are performed or cell volume reaches 1.5 of the initial volume. All simulations described in this section used the turgor pressure *P* = 0.2 MPa. A snapshot of a typical bud emergence simulation is shown in **Figure 5.**

**Figure 5.**
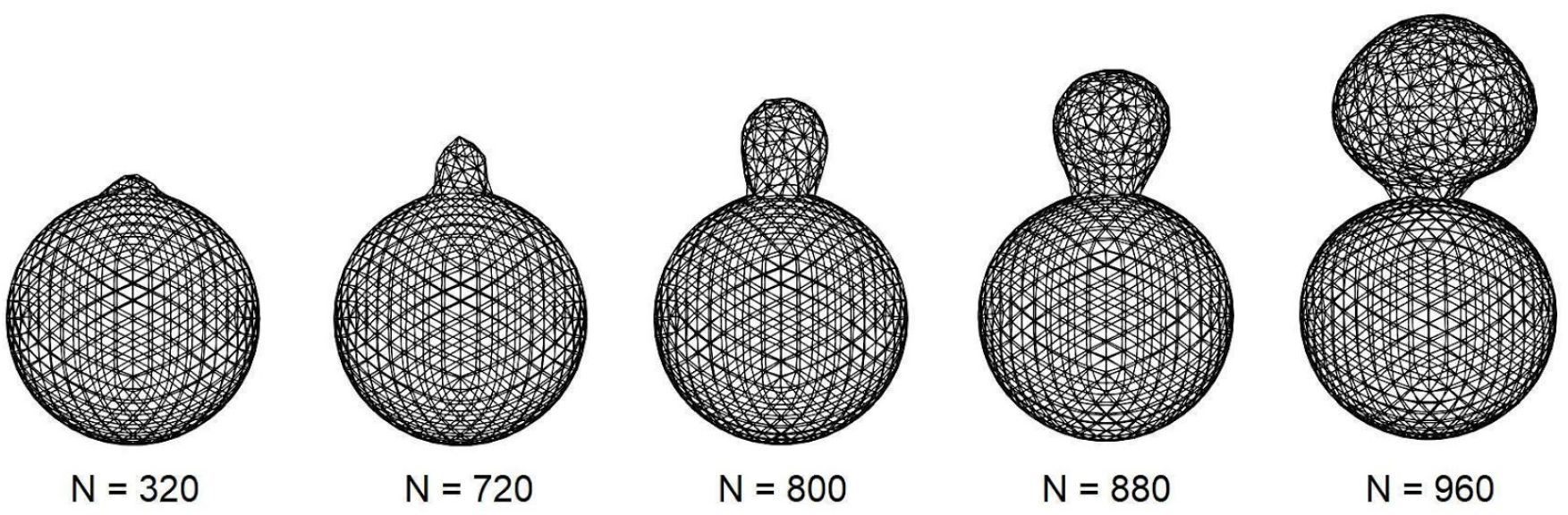
Sample simulation of bud formation under uniformly altered mechanical properties of the cell surface in the budding region. After protrusion occurs, the growth of the bud starts as a tubule growth (left) then transitions to a more spherical expansion (right). The strong constraint from the chitin and septin ring we impose naturally restricts the cell surface expansion at the bud neck. *N* is the number of iterative steps/1000. The relaxation step is Δ*t* = 0.001.

### 3.1 Role of elasticity of cell surface in yeast budding

In this section we keep the weights constant throughout the bud site in time by assuming that polarization of Cdc42 is established before the mechanical properties of the bud are changed. First, the dimensionless ratio of stretching to bending moduli and the critical value 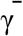 required for bud emergence are determined. Next, their impact on the shape after budding occurs is studied.

#### 3.1.1 Dimensionless ratio of stretching to bending stiffness

In the theory of elasticity, the ratio between the stretching and bending moduli determines the physical property of the material [63,64]. This ratio is often described by the dimensionless Föppl-von-Kármán (*FvK*) number, *k_s_L*_0_^2^/*k_b_*, where *k_s_* is the stretching stiffness, *L*_0_ is an equilibrium edge length of linear spring in the mesh, and *k_b_* is the bending stiffness. By following the same definition, we determine the range of the *FvK* number characterizing the budding region resulting in bud generation.

#### 3.1.2 Bud emergence dependence on the Föppl-von-Kármán number

Simulations with different *FvK* numbers were performed to test whether abud can be generated (**Table 1**). In this section, 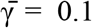 and α_*a*_ = 0.1 are fixed.

**Table 1.** Bud emergence for different *FvK* numbers

| $\alpha_s$ | $\alpha_b$ | $FvK$ number | Bud Emergence |
| --- | --- | --- | --- |
| 0.2 | 0.02 | 3.624 | YES |
| 0.75 | 0.075 | 3.624 | YES |
| 0.2 | 0.05 | 1.450 | NO |
| 0.4 | 0.1 | 1.450 | NO |

The simulations suggest that bud emergence strongly depends on the *FvK* number. Bud emergence does not change if the *FvK* number remains the same. Budding is more likely to occur with a sufficiently large *FvK* number, which can be achieved by either increasing the stretching stiffness or reducing the bending stiffness.

#### 3.1.3 Bud emergence depends on critical elasticity

We first vary bending stiffness at the bud site using weight constant α_*b*_, and the critical value, 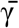, for **Eq. 6,** to study the effect on bud formation for cell surfaces with different fixed *FvK* numbers. Notice that according to the definition, larger α_*b*_ indicates stronger resistance of the cell surface to bending deformation. For each parameter set 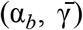, three simulations are performed for the purpose of statistical significance. To qualify as bud emergence, a visible protrusion from the cell surface along with approximately at least 5.5% in total cell volume must be present. If a simulated cell fails the described requirements for bud emergence in all three simulations, it is marked as “No Bud” (**Figure 6**). Different values of α_*b*_ and 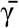 are chosen as described in **Table S2**. The simulation results indicate a clear cutoff value dependent on α_*b*_ for bud emergence and this cutoff reduces as 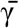 increases (**Figure 6A**).

**Figure 6.**
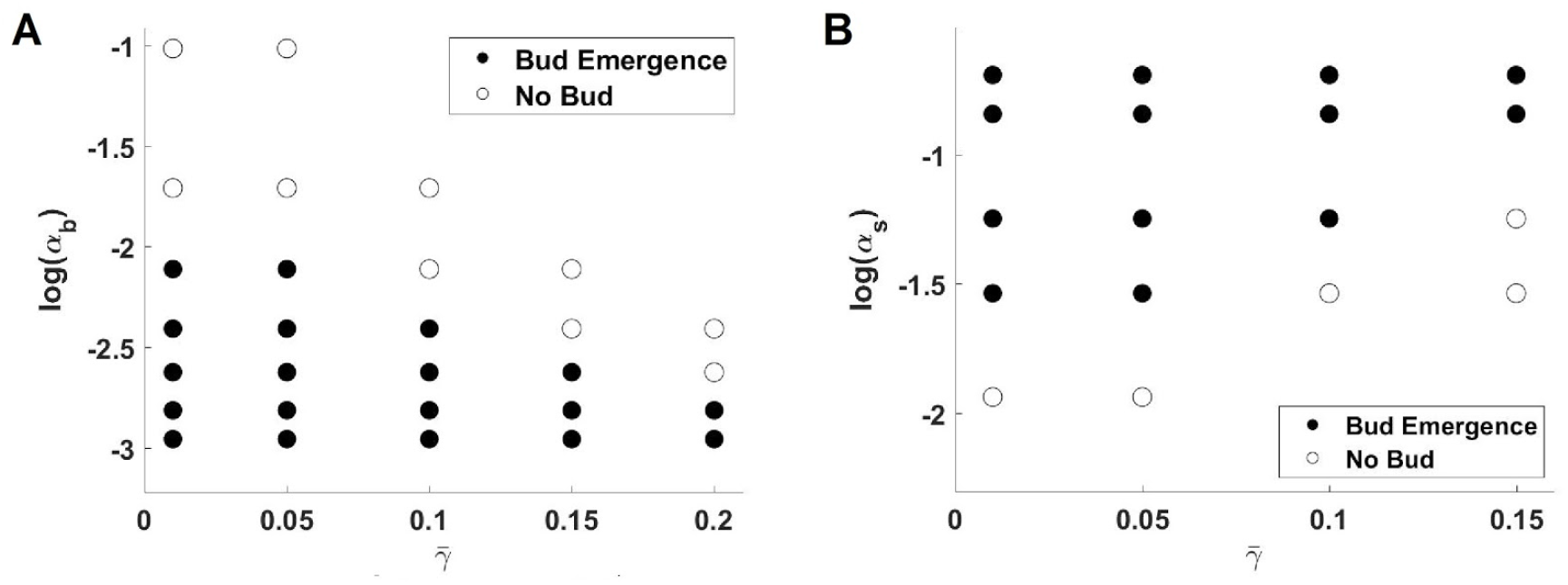
Diagrams describing the influence of 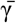 on bud emergence. (A) Variation of bending stiffness (α_*b*_) with fixed α_*s*_ and α_*a*_. (B) Variation of stretching stiffness (α_*s*_) with fixed α_*b*_ and α_*a*_. Both plots show that budding can occur when increasing *FvK* number (dimensionless stretch-bend ratio) over a certain cutoff value, i.e., reducing α_*b*_ or increasing α_*s*_.

Next, simulations were performed with fixed α_*a*_ and α_*b*_, and perturbed α_*s*_ and 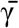. A cutoff value, α_*s*_, for bud emergence was also observed for each value of 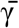 and this cutoff value increases as 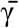 increases (**Figure 6B**). This is expected because bud emergence depends on the physical properties described by the *FvK* number. The same ratio can be achieved by increasing the stretching stiffness or reducing the bending stiffness., while fixing the other.

#### 3.1.4 Impact of the Föppl-von-Kármán number on bud shape

In this section we investigate how different ratios of stretching and bending stiffness affect the bud shape. Since increasing the stretching stiffness is equivalent to reducing the bending stiffness when changing the Föppl-von-Kármán number, in this section we fix α_*s*_, α_*a*_ and vary both α_*b*_ and 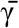. To evaluate the sphericity of the bud, we define 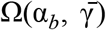 as the distances between the cell surface nodes within the bud area to the center of the bud, for given α_*b*_ and 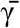 that can generate a bud. Here the bud center is determined by the average of the x-, y-, and z-coordinate of all nodes in the bud region. Therefore, smaller range of 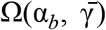 indicates more spherical shape and the average of 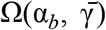 represents the radius of the sphere that fits the bud. We observe that the range of 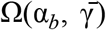 becomes smaller as the weight α_*b*_ applied to the bending modulus of the bud increases for 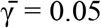 (**Figure 7**). For different 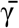 values, we observe the same trend regarding the deviation of 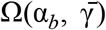 versus α_*b*_ (**Table S3**). This behavior is expected, as higher α_*b*_ leads to stronger resistance to bending deformation at the bud site and therefore more spherical shape can be maintained. We also expect more spherical shapes can be obtained when reducing the stretching modulus.

**Figure 7.**
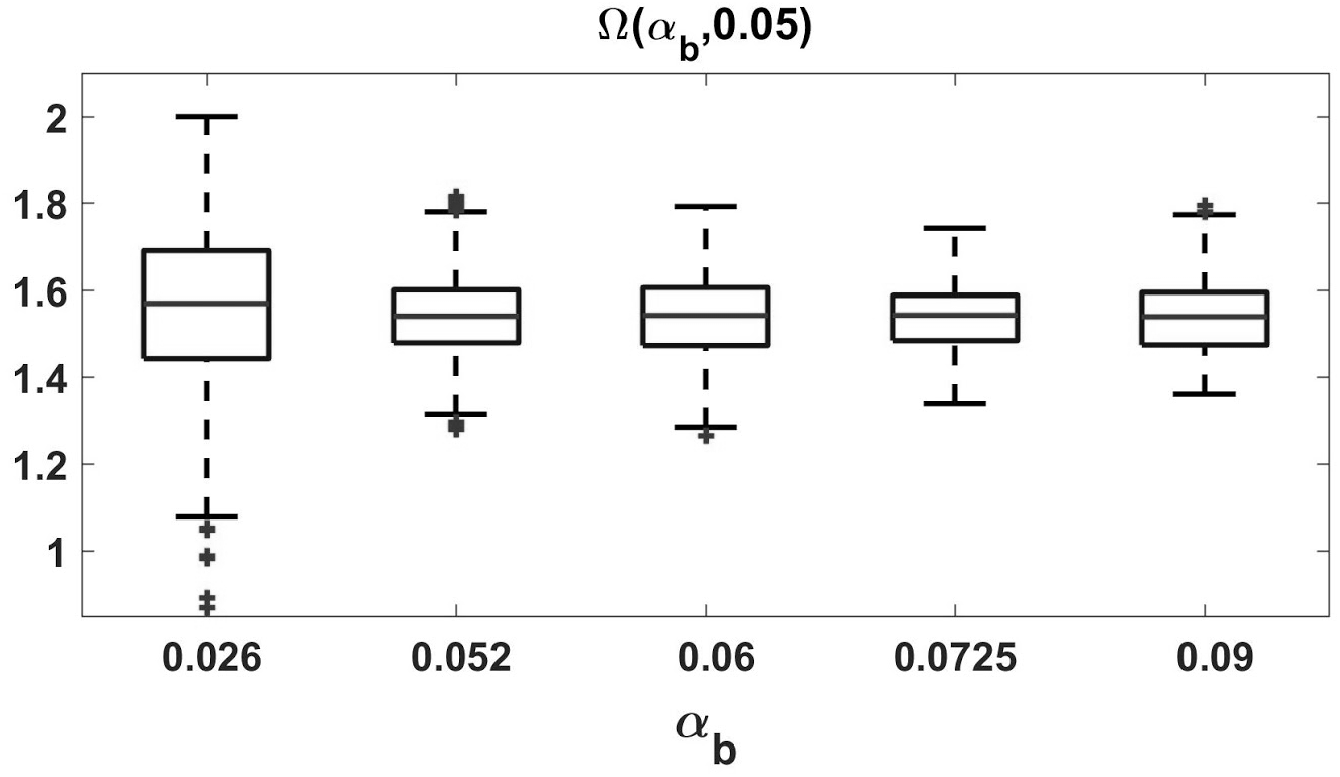
Boxplot of Ω(α_*b*_, 0.05) for different α_*b*_. As α_*b*_ increases, it corresponds to improved resistance to bending in the budding region, hence leading to a narrower range of Ω(α_*b*_, 0.05). The ends of each box denote the 25th and 75th quantiles, denoted *q*_25_ and *q*_75_. Whiskers denote the most extreme data points within 1.5 times the range between *q*_25_ and *q*_75_. Data outside the range are denoted as outliers and shown using black ‘+’s.

Taken together, results in this section suggest a tradeoff based on *FvK* numbers for any fixed critical value for γ, i.e. the dimensionless stretching-to-bending ratio must be sufficiently high at the bud site to generate a bud, while a higher ratio leads to less spherical bud shapes. Therefore, the ratio of stretching to bending moduli must be appropriately tuned by the polarizing molecules to give rise to buds with spherical shapes as observed in experiments.

### 3.2 Role of the budding neck in bud formation

It has been shown in experiments that chitin and septin ring assemblies are important in determining budding neck shape and other growth related activities during a cell cycle [21]. In wild type yeast budding, the bud neck is roughly 1.0 μ*m* in diameter [19], while mutants with impaired chitin and septin ring can exhibit bud necks with approximately 2.68 μ*m* in diameter.

It has also been shown that the septin based ring structure acts as a diffusion barrier to the polarity factors including Cdc42 and cortical proteins, and further affects the bud shape [65]. Here we investigate the mechanical contribution of the chitin and septin rings to bud formation, as well as the shape and size of the bud neck.

Different levels of the rigidity of the combined chitin and septin ring, modeled as an elastic ring with different linear spring coefficients, *k_s_^ring^*, are tested and the bud neck diameter is approximated from each simulation (**Figure 8A**). Consistent with experiments, a direct effect of the decreased constraint of the chitin and septin ring in the simulations is the widened budding neck. More precisely, we observe an exponential decay in the approximated budding neck diameter as *k_s_^ring^* increases (**Figure 8A**).

**Figure 8.**
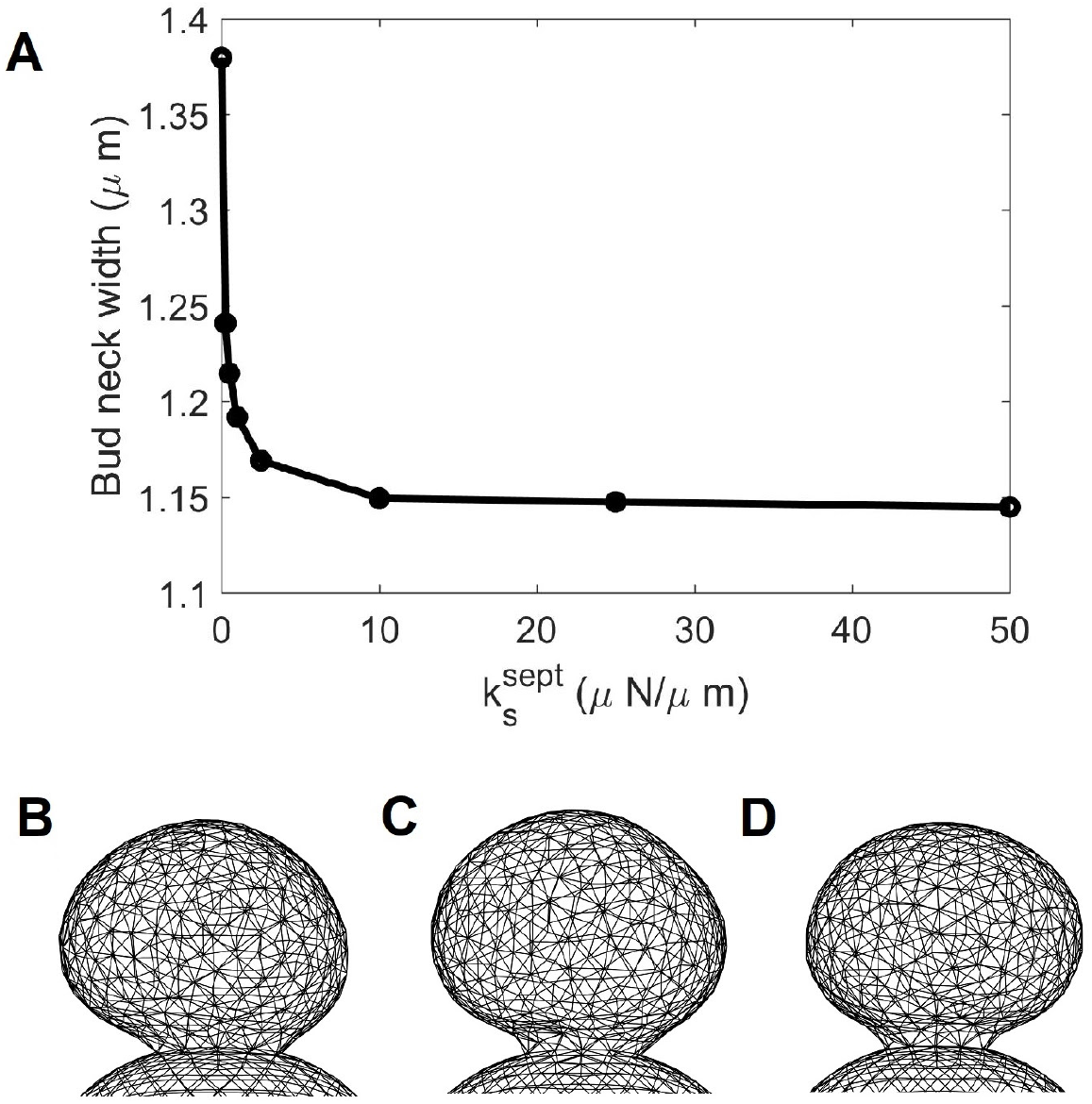
(A) The approximated diameter of the bud neck plotted against stiffness of the ring *k_s_^ring^* = 0,0.25,0.5,1, 2.5,10,25, 50. A sharp decay is observed when *k_s_^sept^* is small. (B - D) Comparison of the bud shape and budding neck with (α_*s*_, α_*b*_, α_*a*_) = (0.5, 0.06, 0.1) fixed and different rigidities of the chitin and septin ring. (B) *k_s_^rms^* = 0.0, (C) *k_s_^rms^* = 0.25, (D) *k_s_^ring^* = 1.0. The corresponding approximated standard deviations are 0.1135, 0.0980, 0.1113 respectively, which are close to the standard deviation 0.0930 with *k_s_^sept^* = 50.0.

Moreover, the standard deviation of the bud radii does not significantly depend on *k_s_^ring^*, ranging between 0.093 and 0.114, indicating that the rigidity of the chitin and septin ring does not impact the bud shape once budding occurs (**Figure 8B-D**). In addition, increase in *k_s_^ring^* can slow down bud growth, or even prevent budding when the *FvK* number is not sufficiently high, as discussed in **SI.2**.

To summarize, in addition to preventing the diffusion of polarity molecules involved in the budding process, the rigid chitin and septin rings impact bud emergence and determine the neck shape without changing the bud shape. The bud neck width reduces as the ring stiffness increases and high rigidity may prevent bud formation.

### 3.3 Bud formation under different polarization patterns

Cdc42 orients actin cables to recruit more Cdc42 as well as new cell membrane and cell wall materials before bud formation. The mechanical properties at the bud site are regulated while the polarization of Cdc42 is established. Budding can start before Cdc42 obtains sharp polarity. In this section, instead of assuming the mechanical properties are regulated by Cdc42 with a steep polarized distribution and using the constant weights (α_*s*_, α_*b*_, α_*a*_) at the bud site, we allow the mechanical properties to undergo a smooth monotonic change from the mother cell to the bud region, which are altered most at the apical tip of the bud site, and study the corresponding conditions for bud emergence and the effect on bud shape.

In particular, we apply Hill functions to model the changes in mechanical properties due to the concentrated Cdc42 along the cell membrane near the bud site, as described in the supplemental information (**SI.3**). All weight functions have maximum value 1.0 in the mother cell, indicating no change in mechanical properties, and the minimum are set to be 0.5 for α_*s*_, 0.052 for α_*b*_, 0.1 for α_*a*_ at the bud tip, and 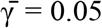. Different Hill coefficients *n* are adopted to model different polarization patterns. Larger *n* indicates a more concentrated change, i.e. the weight functions converge to the constant weights in the previous section as *n* approaches to infinity. We test *n* = 8,17,35,70 to ensure the shapes of the corresponding weight functions are distinguishable in terms of sharpness (**Figure 9A**).

**Figure 9.**
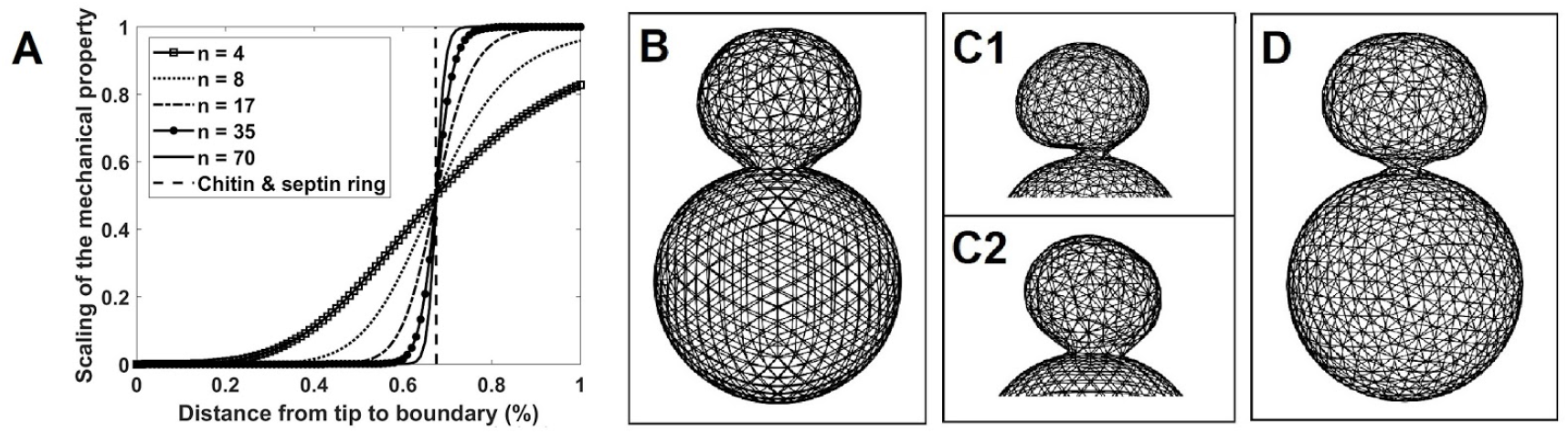
(A) Hill functions with different Hill coefficients used for spatially dependent changes in mechanical properties at the bud site. The Hill coefficients are chosen to be *n* = 8,17,35,70. The weight represents level of the change, with being altered most at distance 0.0 and unaltered at distance 1.0. (B-D) Sample budding shapes based on different Hill coefficients. (B) *n* = 70, (C1, C2) *n* = 35, (D) *n* = 17. Two different shapes of the budding neck are observed for *n* = 35: non-axisymmetric bud neck (C1) and axisymmetric bud neck (C2). Between these two modes, the non-axisymmetric bud neck appears more frequently in simulations.

We found that, for *n* = 8, the cell cannot bud. For *n* = 17, a bud successfully emerges but exhibits a much narrower hourglass-shaped neck with an approximate diameter of 0.4-0.552 μ*m* (**Figure 9D**). For *n* = 35, two different types of budding in terms of the neck shape are observed in simulations: narrower and non-axisymmetric neck (**Figure 9C1**), and similar neck shape as the constant weight cases (**Figure 9C2**). The non-axisymmetric neck shape occurs more frequently in simulations, and the neck diameter ranges 0.87-1.005 μ*m* approximately, which is calculated via the average of the max and min diameters in this case. For *n* = 70, budding can occur in a similar way as was observed when the weights were constant (**Figure 9B**), which is expected for large *n*. Moreover the resulting neck diameter was approximately 1.087-1.14 μ*m*, which is similar to experimental observations.

Overall, the simulations suggest that budding emergence depends on the concentration distribution of the mechanical regulating molecules. The change in the mechanical properties at the bud site controlled by a more polarized signaling molecule is more likely to generate a bud with more robustness in the bud shape. The bud neck obtained with less polarized weight functions becomes narrower and non-axisymmetric.

### 3.4 Bud formation under dynamic change in mechanical properties

Experiments show that Cdc42 polarizes at the apical tip region before bud formation and this highly concentrated distribution is maintained at the early stage of bud growth. As formation takes place, Cdc42 aggregates to reach a homogeneous distribution within the bud site [6]. Before cell division, Cdc42 is redirected from the bud cortex to the bud neck. This suggests that regulation of the mechanical properties might change temporally with strongest effect before bud formation or right after the apical protrusion. Therefore, we test a temporal restoration of altered mechanical properties at the bud site in our model to see whether the strong mechanical regulation, if only present in a short period at the beginning of the budding process, is sufficient or not.

#### 3.4.1 Temporal restoration functions

Due to the energy dissipation approach used for mechanical relaxation in our simulations, the temporal restoration of the bud mechanical properties is assumed to be cell volume based, i.e., the evolution in time of the weights in altering mechanical potentials at the bud site is assumed to be linearly increasing with respect to the volume:

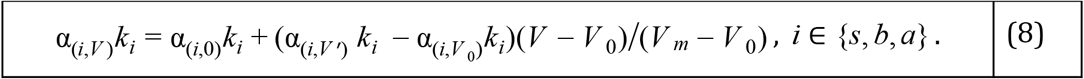

Here α_(*i,V*)_ represents the weight applied to the linear spring potential (*k_s_*), cosine bending potential (*k_b_*), or the area expansion resistance (*k_a_*), based on the current volume *V*. When the volume reaches the target volume *V_m_*, the mechanical properties of the bud become identical to those of the mother cell. Here, *V*_0_ represents the initial volume of the cell.

#### 3.4.2 Temporal restoration of bud mechanical properties leads to symmetric bud shape

To test different restoration speeds from the altered state, we change the value of *V_m_*. For example, the mechanical properties at the bud site will be fully restored to the same level as the mother cell when the cell volume doubles, i.e. *V_m_* = 2*V*_0_.

Similarly, setting *V_m_* = 1.5*V*_0_ and *V_m_* = 3*V*_0_ lead to expedited and delayed restoration compared to *V_m_* = 2*V*_0_, respectively. Based on the results of model calibration, we test the following parameter set as the altered state: (α_*s,V*_0__ α_*b,V*_0__, α_*a,V*_0__, 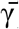) = (0.5, 0.0151, 0.1, 0.05). In simulations, the weights α_*s*_, α_*b*_, α*_a_* are changed in time following Eq. (8). We found that temporal restoration of the mechanical properties leads to bud formation with more spherical and symmetric shapes (**Figure S3**). Regardless of different choices of the restoration speed, improved bud roundness was observed once the bud formed (**Figure 10A**). Namely, we compared the local standard deviations between buds with total cell volume of 1.39*V*_0_ – 1.45*V*_0_ and 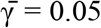, which give rise to buds in asymmetric growth when applying the constant uniformly altered mechanical properties at the bud site. This is quantified using local standard deviations (SD) include upper hemisphere, *SD_u_*, lower hemisphere, *SD_l_*, and SD away from the neck, *SD_n_. SD_u_* is calculated using cell surface nodes positioned above the center of the bud. *SD_l_* is calculated using cell surface nodes positioned between the bud neck and the center of the bud. *SD_n_* is calculated by considering only nodes whose z-coordinates are in the upper 90% of the height of the bud. The optimal values, based on *SD_n_*, are presented in **Table 2.**

**Table 2:** Optimal local SD of the bud for uniformly altered and restorative mechanical properties

| | Overall SD | $SD_u$ | $SD_l$ | $SD_n$ |
| --- | --- | --- | --- | --- |
| $\alpha_b = 0.052$ | 0.0516 | 0.0354 | 0.0454 | 0.0423 |
| $\alpha_b = 0.06$ | 0.0840 | 0.0683 | 0.0943 | 0.0753 |
| $\alpha_b = 0.09$ | 0.0796 | 0.0695 | 0.0810 | 0.0756 |
| $\alpha_b = 0.121$ | 0.0514 | 0.0409 | 0.0596 | 0.0425 |
| $V_m = 2.0V_0$ | 0.0415 | 0.0368 | 0.0455 | 0.0350 |
| $V_m = 3.0V_0$ | 0.0568 | 0.0544 | 0.0577 | 0.0552 |
| $V_m = 1.75V_0$ | 0.0369 | 0.0282 | 0.0438 | 0.0268 |

**Figure 10.**
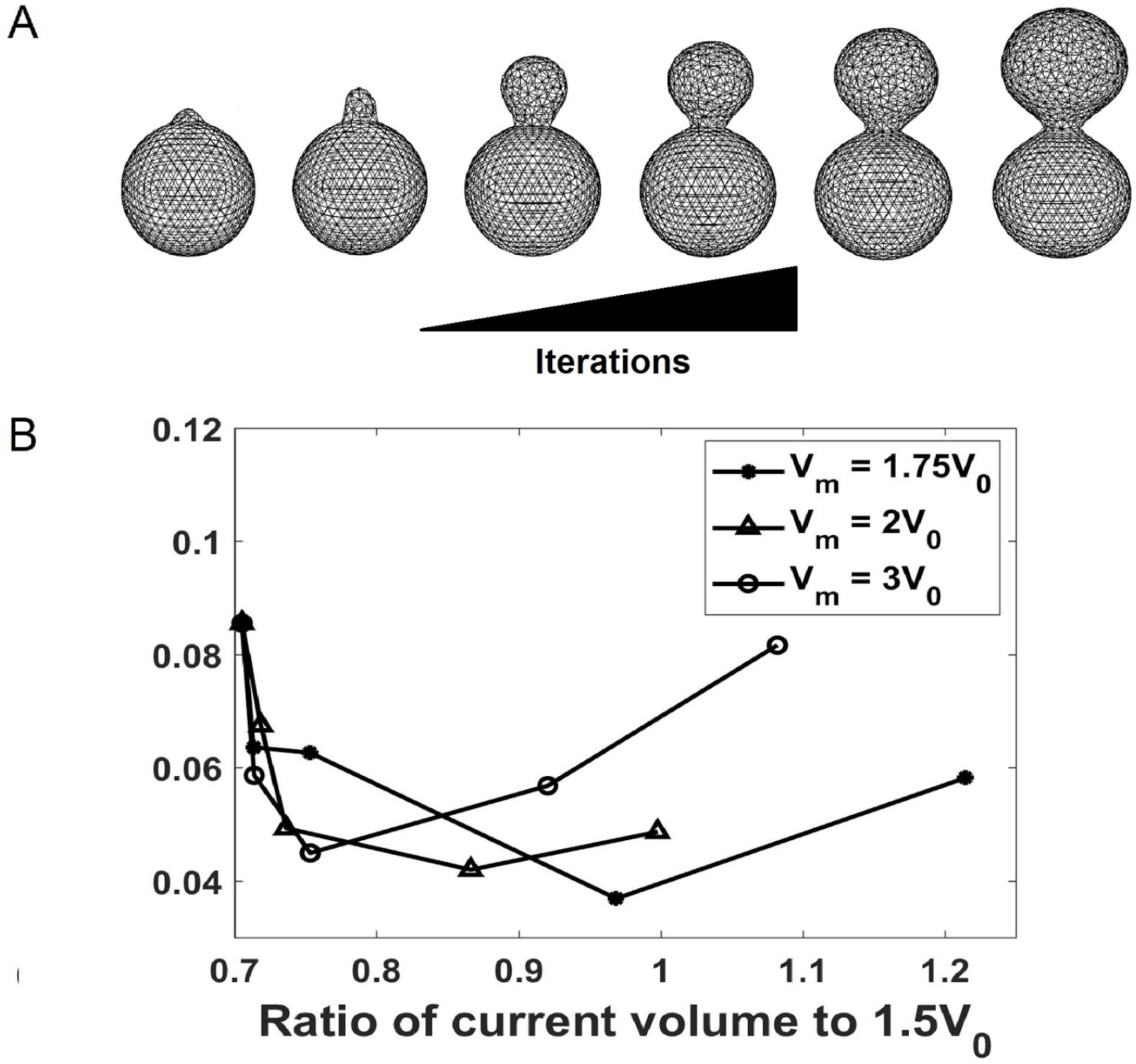
(A) Sample simulation of bud formation under temporal restoration of the cell surface mechanical properties in the budding region. After protrusion occurs, the growth of the bud starts as a tubule growth (left) then transitions to a more spherical expansion (right) compared to the simulation with time-independent changes in mechanical properties (**Figure 5**). In this example, *V_m_* = 2*V*_0_. (B) Standard deviation of bud radii vs. ratio of current volume to 1.5 *V*_0_. Note that higher target volume implies slower restoration speed, which exhibits faster transition from apical growth to isotropic growth, and later transitions into asymmetric growth as observed in the time-independent change in mechanical properties cases.

In addition, we observed that the restoration speed had a significant impact on bud formation, as faster restoration led to bud inhibition, such as *V_m_* = 1.5 *V*_0_. On the other hand, when the restoration speed was slower, such as *V_m_* = 3.0*V*_0_, the simulation showed asymmetric growth.

The standard deviation of radii at the bud region are calculated for cases with *V_m_* = 1.75*V*_0_, 2*V*_0_, 3*V*_0_ (**Figure 10**). In particular, for *V_m_* = 3*V*_0_, the bud is initially generated in a spherical shape. Due to the slow restoration of mechanical properties allowing efficient expansion of the bud, the bud shape gradually becomes more ellipsoidal, thereby achieving a standard deviation of bud radii similar to which is observed under time-independent changes in mechanical properties (**Table S3**). This suggests a well-coordinated restoration of the bud mechanical properties may be necessary for symmetry maintenance of the bud. For *V_m_* = 2*V*_0_, a more symmetric bud shape is maintained after protrusion in comparison with the time-independent cases presented in Section 3.1 (**Figure 10A**).

## 4 Discussion and conclusions

In this paper, a novel 3D coarse-grained particle-based model of a single cell is introduced and used to investigate the role of local changes in cell surface mechanical properties during the early stages of yeast budding. The model combines nonhomogeneous representation of the cell surface with stiff ring structures to study cell growth and budding. It is calibrated using experimentally measured Young’s modulus of the budding yeast cell wall [41]. The novelty of this study lies in testing a novel hypothesised mechanism of cell budding combining changing mechanical properties of the budding region with the impact from the constraint of chitin and septin rings.

By assuming mechanical properties are altered uniformly at the bud site by the polarized molecules, bud emergence was shown to depend on the Föppl-von-Kármán (*FvK*) number (Section 3.1). Budding can occur only when the stretching modulus is large or the bending modulus is sufficiently reduced. As the value of 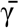 increases, bud formation requires a higher *FvK* number. One explanation for the requirement of the altered ratio at the bud site for bud emergence is that a reduced bending stiffness or increased stretching stiffness can facilitate bending deformation and thus promote bud formation.

For an emerging bud, the shape symmetry is biologically important because it indicates the balance between the composition and integrity of the bud surface, which is observed in wild type yeast budding. The sphericity of the bud shape, described via standard deviation of the bud radii, was shown to become lower as the *FvK* number was reduced by the polarized molecules or the critical value of *γ* (Eq. 6) increased in simulations.

It is known that the chitin and septin rings serve as diffusion barriers for polarity factors at the early stage of budding. By using a computational model, we found that the resistance provided by the stiff rings prevents or slows bud growth. These ring structures also determine bud neck shape in a way that the neck diameter reduces as the stiffness increases, without affecting the bud shape.

By assuming mechanical properties are altered by polarizing molecules with different polarization distributions at the bud site, it was shown that one cell was more likely to generate a bud with a more symmetric shape under a more polarized distribution. The bud neck obtained with less polarized distribution was narrower and non-axisymmetric.

By incorporating a dynamical restoration of mechanical properties at the bud site, the resulting bud shape was shown to be more symmetric and spherical, as compared with the ones with time-independent changes in mechanical properties. Fast restoration prevents bud formation and slow restoration leads to development of an asymmetric bud.

The 3D model described in this paper, although calibrated by using data for budding yeast, can be applied to study cell growth and budding in other biological systems. It allows one to study the effect of local regulation of mechanical properties leading to global morphological changes. For instance, investigating how adverse effects due to local perturbations in the mechanical properties can propagate during growth is important for getting a better understanding of morphogenesis. In future, we plan to include in the modeling approach dynamic biochemical signaling networks submodel coupled with the cellular and subcellular mechanical submodels.

Namely, to investigate the regulation of the mechanical properties more precisely, it is worthwhile to include the model of spatiotemporal dynamics of the signaling molecules. The triangular mesh used for the mechanical model, with re-meshing techniques implemented to improve the element regularity, would be used to simulate diffusion-reaction systems of equations describing biochemical signals. Several studies have already studied the connection between biochemical signalling and cell mechanics in mating yeast and tubule formation [39,66,67]. However, understanding of the coupling between the biochemical signaling pathways and cell mechanics for the yeast budding process is still incomplete. A biologically calibrated mechano-chemical model would be helpful providing predictions to be tested in future experiments.

The model described in this paper oversimplifies contributions from cytosol components of the cell including nucleus, vacuole and actin-filaments. At the same time, the actin-filaments play a very significant role in cell surface expansion [68–70]. The nucleus also plays an important role especially in G2 and M phases of the cell cycle [71]. Recently our group developed a more refined 2D mechanical SCE type model with detailed description of the nucleus, actomyosin and cadherin, to study tissue bending mechanisms in a wing of a Drosophila embryo [72]. We are now working on incorporating these new components as well as biochemical submodel into our 3D model of an asymmetric cell growth which can be also used for studying viral budding and other vegetative reproduction processes performed via budding.

## Acknowledgements

The authors acknowledge partial support from the National Science Foundation Grant DMS-1762063 through the joint NSF DMS/NIH NIGMS Initiative to Support Research at the Interface of the Biological and Mathematical Sciences and the National Science Foundation Grant DMS-1853701. RZ was supported by the National Science Foundation through Grant No. DMR-1719550. Simulations were performed using the computer clusters and data storage resources of the HPCC at University of California, Riverside, which were funded by grants from NSF (MRI-1429826) and NIH (1S100D016290-01A1)

