## Supplemental Information for "Role of combined cell membrane and wall mechanical properties regulated by polarity signals in cell budding"

### SI.1 Effect of the bending stiffness on the evolution of the bud shape

The target cell volume, $1.5V_0$, is a termination criterion for simulations. We record and analyze the standard deviation of the budding process using simulations described in Sections 3.1.2 and 3.1.3 that reached the target volume. Based on the simulation results, we observe an overall downward trend of the standard deviation at the early- to mid-stage of the simulation, which corresponds to the transition from apical growth to isotropic growth. Qualitatively this trend is represented by the first four snapshots of the sample simulations (**Figure 5**). Before reaching the target volume, a majority of the standard deviations increase, and cases such as $\Omega(0.052,\ 0.01)$ and $\Omega(0.0725,\ 0.01)$ show a sharp increase (**SI Figure 1**). However, this increase in the standard deviation can be biased due to the inclusion of the budding neck into the calculation of $\Omega(\alpha_b, \bar{\gamma})$. The increase in standard deviation can be alleviated by increasing the weight for bending stiffness or by increasing the critical value of $\gamma$. Out of all trajectories, only $\Omega(0.52,\ 0.2)$ shows consistent decrease in the standard deviation. $\Omega(0.6,\ 0.2)$, on the other hand, shows a jump in the standard deviation when the cell volume approaches the target despite higher $\alpha_b$, suggesting the behavior of $\Omega(0.52,\ 0.2)$ may be an isolated case.

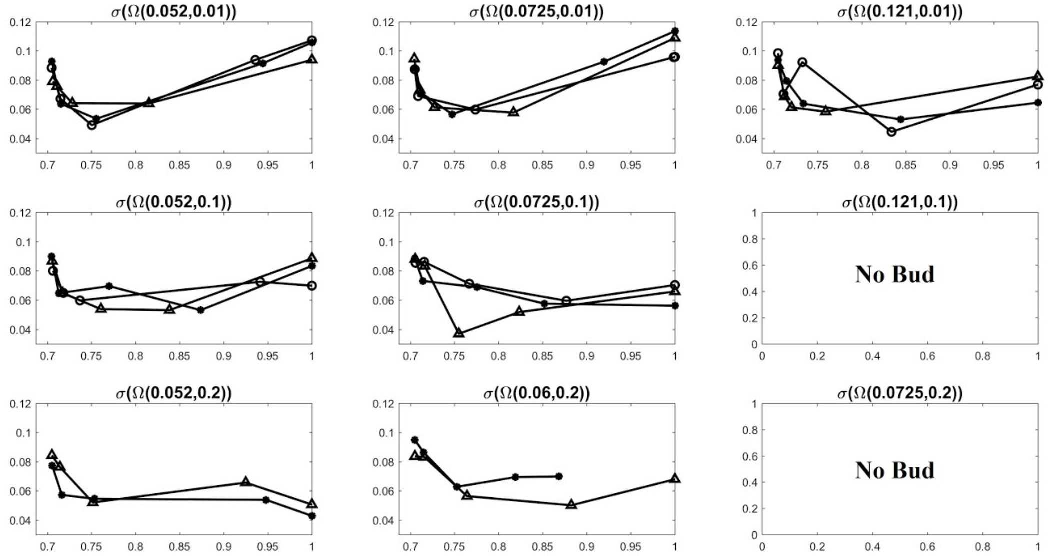


**SI.Figure 1.** Trend of standard deviation of selected $\Omega(\alpha_b, \gamma)$. The x-axis in each plot represents the ratio of current cell volume to the target cell volume, $1.5V_0$. The standard deviation for buds with the volume close to the target value is increased more when both $\alpha_b$ and $\bar{\gamma}$ are small.

### SI.2 Stiff chitin and septin rings slow down or prevent budding

To demonstrate the impact of bud neck constraint originated from the chitin and septin ring on bud emergence, we choose parameter sets where bud emergence takes extraordinarily long time or is inhibited under strong bud neck constraint. For $(\alpha_s, \alpha_b, \alpha_a, \bar{\gamma}) = (0.5, 0.121, 0.1, 0.05)$, buds can be formed with $k_s^{ring} = 50.0$ and $k_s^{sept} = 1.0$ but the former takes about $4.12 \times 10^6$ iterative steps to reach the bud size which is half of the mother cell, while the latter takes roughly $1.06 \times 10^6$ iterative steps. Furthermore, we let $(\alpha_s, \alpha_b, \alpha_a, \bar{\gamma}) = (0.5, 0.181, 0.1, 0.05)$ and test the same $k_s^{ring}$ values, and find bud inhibition in both $k_s^{sept}$ values. This shows that strong bud neck constraint from the chitin and septin ring can be challenging for bud emergence if the Föppl-von-Kármán number is not sufficiently high.

### SI.3 Using Hill function to model the spatially dependent weights

A typical Hill function can be formulated as $f(x) = min + (max - min)/(1 + \left(\frac{x}{K}\right)^n)$ where $x$ denotes the spatial position, $K$ is a constant indicating the position corresponding to the midpoint between min and max, i.e., $f(K) = 0.5(min + max)$. $n$ is the Hill coefficient that determines the sharpness of the gradient changing from min to max, causing the function shape to be linear when $n = 1$ and change to stepwise when increasing $n$ (**Figure 8**). Because of this shape variation, we model Cdc42 distributions at different stages of polarization using a Hill function with different values of $n$.. We define the weight as a spatial function of Hill type $\alpha(x) = 1/(1 + \left(\frac{x}{K}\right)^n)$ for each mechanical potential coefficient, such that $\alpha(x)$ is between 0 and 1. The Föppl-von-Kármán number is not altered when multiplied by weight 1, corresponding to the region of the mother cell, while the it is maximized when multiplied by weight 0, corresponding to the apical tip of the bud site.

To couple this Hill type weight function in the model, we start by selecting a reference point $X_0$, which is determined via the average of x- and y-coordinate of all cell surface nodes, and the z-coordinate of the tip of the budding region. The location of the chitin and septin rings relative to the average radius of the bud is always set to be the midpoint in the Hill function, $K$, in order to study the effect of sharpness of the changes in mechanical properties. Moreover, $K$ is updated as the bud grows, such that the relative location of the midpoint of the weight function remains the same throughout the simulation.

#### SI.4 Role of chitin and septin rings by using Hill function to model the spatially dependent weights

We perform the same model simulations with spatially dependent weight functions by altering the mechanical properties at the bud site as described in Section 3.3. Simulations performed with the Hill type weight functions are in agreement with the cases with constant weights. Specifically, the bud neck width reduces rapidly when $k_s^{ring}$ increases from 0 to 1 and remains more or less the same when $k_s^{ring}$ keeps increasing to 50 (**SI Figure 2**).

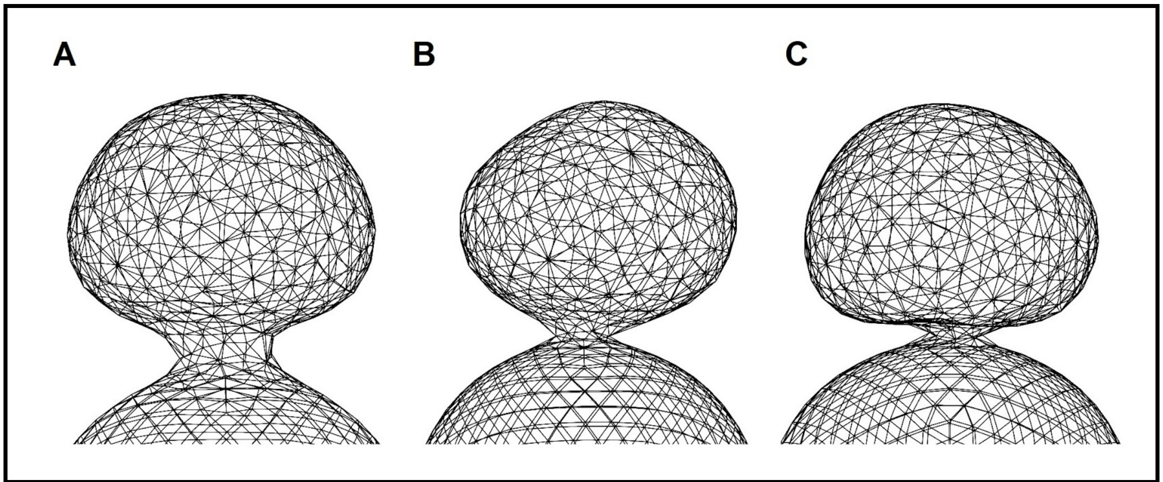


**SI Figure 2.** Comparison of the bud shape and budding neck under nonuniform, spatial-dependent changes in mechanical properties with different rigidities of the chitin and septin ring. The Hill coefficient is 17 and (A) $k_s^{ring} = 0.0$, (B) $k_s^{ring} = 1.0$, (C) $k_s^{ring} = 50.0$. In all three cases, the budding is initiated near the center of the budding region away from the chitin and septin ring location. However, simulations with weaker bud neck constraint lead to wider budding necks.

### SI.5 Comparison of bud shapes obtained by time-independent change in bud mechanical properties vs. temporally restored mechanical properties

We compare the bud shapes with the similar volumes under different patterns of changes in the mechanical properties. According to the standard deviation of bud radii, bud shape is more spherical with the temporal restoration as described in **Section 3.4** (**SI Figure 3**).

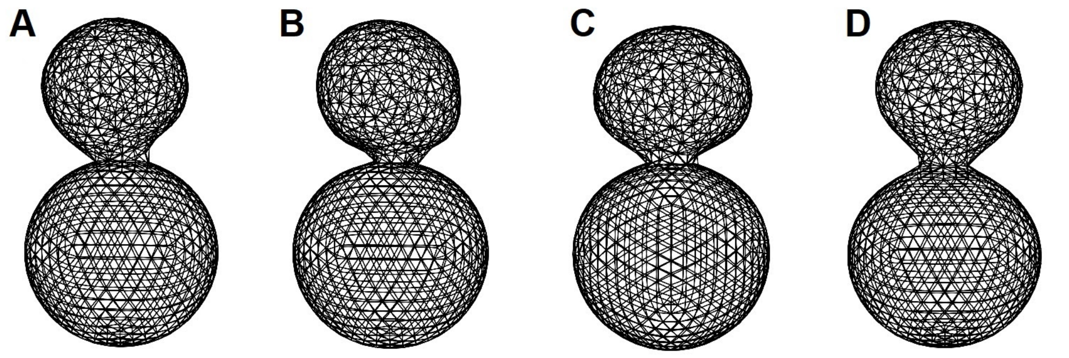


**SI Figure 3.** Comparison between bud shapes obtained using static mechanical properties (**Section 3.1**) and using temporal restorative mechanical properties (**Section 3.4**). Parameter sets used are as follows: (A) $(\alpha_s, \alpha_b, \alpha_a, \bar{\gamma}) = (0.5,\ 0.09,\ 0.1,\ 0.1)$, (B) $(\alpha_s, \alpha_b, \alpha_a, \bar{\gamma}) = (0.5,\ 0.052,\ 0.1,\ 0.05)$, (C) $(\alpha_s, \alpha_b, \alpha_a, \bar{\gamma}) = (0.5,\ 0.121,\ 0.1,\ 0.01)$, (D). Restorative mechanical properties with initial $(\alpha_s, \alpha_b, \alpha_a, \bar{\gamma}) = (0.5,\ 0.0151,\ 0.1,\ 0.05)$. The visual comparison is made at cell volume $V \approx 1.39 - 1.45V_0$.

### SI.5: Tables

#### SI. Table 1: Model parameters of the yeast mother cell

| Parameter | Description | Value(s) | Source |
|---|---|---|---|
| $k_s$ | Linear spring coefficient | $2.0\ \mu N/\mu m$ | Calibration using AFM data [1] |
| $k_b$ | Bending spring coefficient | $0.5\ \mu N \mu m$ | Calibration using AFM data [1] |
| $k_a$ | Area expansion resistance coefficient | $2.0\ \mu N$ | Calibration using AFM data [1] |
| $k_s^{ring}$ | Linear spring coefficient, combined chitin and septin ring | $50.0\ \mu N/\mu m$ | Model assumption based on qualitative observation. |
| $L_0$ | Initial edge length | $0.301\ \mu m$ | Relaxed initial system |

| | | | |
|---|---|---|---|
| $\theta_0$ | Initial dihedral angle | 0.08725 rad | Relaxed initial system |
| $L_0^{rep}$ | Volume exclusion length | 0.301 $\mu m$ | Qualitative observation |
| $D$ | Morse potential well depth | 0.01 | Qualitative observation |
| $a$ | Morse potential well width | 9.0 | Qualitative observation |
| $A_0$ | Initial triangular area | 0.03927 $\mu m^2$ | Relaxed initial system |
| $P$ | Turgor pressure | 0.2 MPa | [2,3] |

**SI.Table 2: Weights and critical values of expansion strain used in section 3.1**

| | | |
|---|---|---|
| Fixed Parameters | $\alpha_s$ | 0.5 |
| | $\alpha_a$ | 0.1 |
| Variables | $\alpha_b$ | 0.052 , 0.060 , 0.0725 , 0.090 , 0.121 , 0.181 , 0.362 |
| | $\bar{\gamma}$ | 0.01 , 0.05 , 0.1 , 0.15 , 0.2 |

**SI. Table 3. Average standard deviation with different $\alpha_b$ and $\gamma$.**

| | $\alpha_b = 0.052$ | $\alpha_b = 0.06$ | $\alpha_b = 0.0725$ | $\alpha_b = 0.09$ | $\alpha_b = 0.121$ |
|---|---|---|---|---|---|
| $\bar{\gamma} = 0.01$ | 0.1005 | 0.1014 | 0.1055 | 0.0746 | 0.0748 |
| $\bar{\gamma} = 0.05$ | 0.0893 | 0.0930 | 0.0743 | 0.0771 | 0.0514 |
| $\bar{\gamma} = 0.10$ | 0.0796 | 0.0699 | 0.0724 | 0.0476 | No bud |
| $\bar{\gamma} = 0.15$ | 0.0586 | 0.0547 | 0.0734 | No bud | No bud |
| $\bar{\gamma} = 0.20$ | 0.0476 | 0.0644 | No bud | No bud | No bud |

Bud emergence is determined via visual inspection such that the protrusion at the cell surface needs to arrive at the level depicted in **Figure 5**, $N$ =720.

**SI. Table 4. Explanation of terminology**

| Terminology | Clarification |
|---|---|
| Area expansion resistance | The harmonic area conservation potential. |
| Time-independent changes in mechanical properties | The assigned mechanical properties remain unchanged in the bud cell surface through the simulation. |
| Spatial-dependent changes in mechanical properties | The assigned mechanical properties change according to the current position on the bud cell surface. |
| Time-dependent changes in mechanical properties | The assigned mechanical properties change according to the progress in the simulation, |
| Föppl-von-Kármán number (dimensionless stretching to bending stiffness ratio) | Describes the ratio between linear spring modulus and bending modulus, scaled by the equilibrium length of the linear spring to ensure a dimensionless number. |
| Strain associated with area expansion | The relative change in area of triangle(s). A threshold value is assigned to determine when new cell surface material is introduced. |
| Dihedral angle | The angle between two intersecting planes. It can be described by the angle between unit normal vectors of the intersecting planes. |
